# Heavy–tailed neuronal connectivity arises from Hebbian self–organization

**DOI:** 10.1101/2022.05.30.494086

**Authors:** Christopher W. Lynn, Caroline M. Holmes, Stephanie E. Palmer

**Author notes:** **Corresponding Author.** Correspondence and requests for materials should be addressed to C.W.L.

## Abstract

In networks of neurons, the connections are heavy–tailed, with a small number of neurons connected much more strongly than the vast majority of pairs.^1–6^ Yet it remains unclear whether, and how, such heavy–tailed connectivity emerges from simple underlying mechanisms. Here we propose a minimal model of synaptic self–organization: connections are pruned at random, and the synaptic strength rearranges under a mixture of Hebbian and random dynamics. Under these generic rules, networks evolve to produce scale–free distributions of connectivity strength, with a power–law exponent 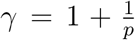 that depends only on the probability *p* of Hebbian (rather than random) growth. By extending our model to include correlations in neuronal activity, we find that clustering—another ubiquitous feature of neuronal networks^6–9^—also emerges naturally. We confirm these predictions in the connectomes of several animals, suggesting that heavy–tailed and clustered connectivity may arise from general principles of self–organization, rather than the biophysical particulars of individual neural systems.

## Main

Neurons communicate and interact via an intricate web of synaptic connections. Although this wiring is constantly shifting and rearranging,^10–12^ its structure is far from random, with synaptic connectivity concentrated among a select few pairs of neurons.6 Indeed, the distribution of connection strengths is heavy–tailed,^1–5^ with a sparse network of strong connections dominating in a sea of weak interactions. These strong connections form the backbone of neuronal circuitry that is responsible for unparalleled feats of information processing,^13^ from learning^10,14–16^ and memory^17–20^ to sensing,^21–25^ communication,^26,27^ and motor control.^28–31^ Yet, despite the importance of strong connections in guiding neural function, there remain fundamental questions about how heavy–tailed connectivity arises in neuronal populations. Does it require complicated biophysical processes specific to each species or neural system? Or, instead, can heavy–tailed connectivity emerge naturally from simple principles of network self–organization?

### Heavy–tailed neuronal connectivity

To answer these questions, we begin by comparing the distributions of connection strengths across several different species. Gathering sufficient data to characterize the tails of connection strength distributions presents a major challenge, requiring cellular–resolution maps of large volumes of nervous tissue.^32^ However, recent advances in high–throughput electron microscopy and image processing have allowed unprecedented snapshots of the physical wiring connecting thousands to tens of thousands of neurons.^1,14,15,21,22,28,29^ Given this experimental progress, it is now possible to investigate the general mechanisms underlying large–scale neuronal connectivity.

Consider, for example, the partial connectome of the *Drosophila* central brain (Fig. 1a), with 14 million synapses linking over 21 thousand neurons (approximately one quarter of the fly’s brain).^1^ Despite the large number of synapses, 99% of neuron pairs remain unconnected; and even among connected neurons, 50% are only linked by a single synapse (Fig. 1b). Meanwhile, some pairs of neurons are connected by over one thousand individual synapses; these rare but extremely strong connections comprise the heavy tail of synaptic connectivity (Fig. 1b). By contrast, if we randomize the placement of synapses in the network, then connectivity strength drops off super– exponentially as a Poisson distribution (Fig. 1b), and the backbone of strong connections vanishes. We observe similarly heavy–tailed connectivity in the *Drosophila* optic medulla (Fig. 1c),^22^ the connectome of the roundworm *C. elegans* (Fig. 1d),^28^ and the sensory–motor circuit of the annelid *Platynereis* (Fig. 1e).^29^

**Fig. 1.**
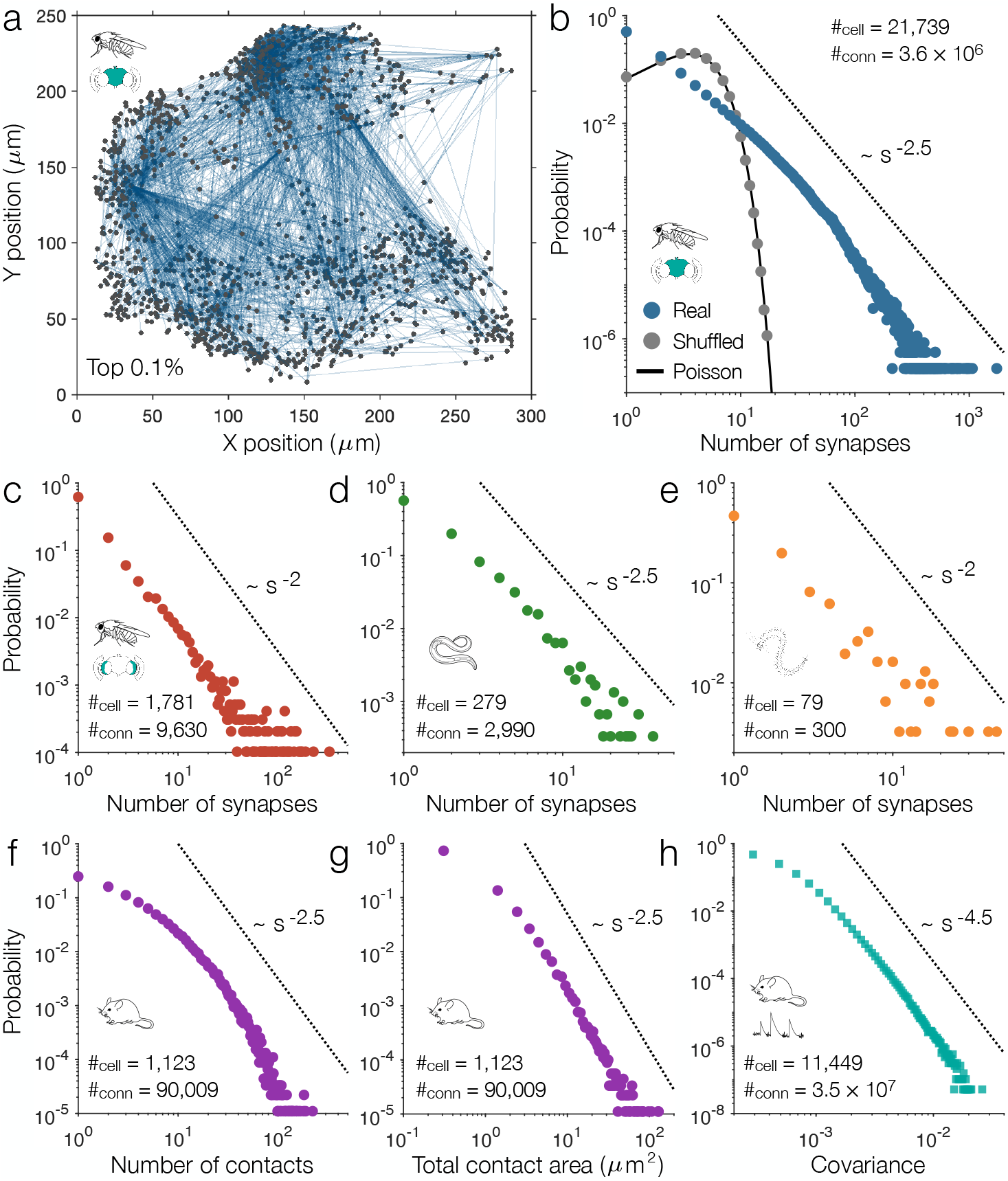
Heavy–tailed connectivity in neuronal connectomes. **a**, Network of the strongest 0.1% of connections in the connectome of the *Drosophila* central brain, traced using volume electron microscopy (see Methods).^1^ **b**, Distribution of the number of synapses in each connection of the *Drosophila* central brain (blue) and after shuffling the synapses among the neurons (grey). The solid line indicates a Poisson distribution, and the dashed line illustrates a power law for comparison. **c-e**, Distributions of the number of synapses in each connection of the *Drosophila* optic medulla (**c**),^22^ the entire connectome of the roundworm *C. elegans* (**d**),^28^ and the sensory–motor circuit of the annelid *Platynereis* (**e**).^29^ **f-g**, Distributions of the number of contacts (**f**) and total contact area (**g**) between neurons in the inner plexiform layer of the mouse retina.^21^ **h**, Distribution of covariances in neuronal activity in the mouse visual cortex while responding to natural images, recorded using two–photon calcium imaging (see Methods).^33^ Square data points are used to differentiate from the structural connectivity in panels **a**-**g**.

Thus far we have focused on the connectomes of invertebrates, in which stronger connections are formed by increasing the number of synapses in parallel.^1^ In vertebrates, meanwhile, connections are strengthened by increasing both the number and size of individual synapses.^1,34^ In the mouse retina, for example, we find that the distribution of the number of physical contacts between neurons (a proxy for synaptic connections) is heavy–tailed (Fig. 1f), but not to the same degree as invertebrate connectivity (Fig. 1b-e).^21^ This accounting, however, fails to include information about the sizes of individual contacts or synapses, which themselves have been observed to follow heavy–tailed (often log–normal) distributions.^5,6,14^ Indeed, if we consider the total strengths of connections—that is, the total contact area between neurons—we find a distribution that once again resembles those observed in invertebrates (Fig. 1g). Finally, leveraging recent large–scale recordings of neuronal activity, we can examine the strengths of “functional” connections (or correlations) between neurons. In a large–scale recording of the mouse visual cortex responding to natural images,33 we find that the distribution of covariances in activity reflects the heavy–tailed structural connectivity between neurons (Fig. 1h), a result that we confirm in different mice and across distinct stimuli (see Supplementary Information).

### Model of Hebbian self–organization

The above results suggest that heavy–tailed connectivity is a general feature of neuronal networks. It is tempting, then, to seek an equally general explanation. In the study of complex systems, heavy-tailed distributions often arise from preferential attachment or rich–get–richer mechanisms. In the context of synaptic connectivity, a similar mechanism already lies at the core of network formation: Hebbian plasticity, wherein strongly–connected neurons are more likely to co–fire, leading to even further growth.^11,34–36^ But can the preferential attachment of connectivity on its own (that is, even before including activity–dependence and other biophysical effects) explain the heavy tails observed in real connectomes? To answer this question, here we present a minimal model of network formation in which connectivity self–organizes under a mixture of Hebbian and random dynamics. Specifically, we begin with *N* neurons connected by *S* units of connection strength (for example, each unit could represent a synapse or 1*μm*^2^ of contact area). The network is defined by a connectivity matrix *A*, where *A_ij_* is the directed connection strength from neuron *i* to neuron *j*, such that ∑_*ij*_ *A_ij_* = *S*. At each point in time, one connection is selected at random to be pruned, resulting in *A_ij_* → 0 (Fig. 2a, left). Each unit of pruned strength is then redistributed to a new pair of neurons (such that *A_ij_* → *A_ij_* + 1) in one of two ways: (i) with probability *p*, a connection is selected for growth in a Hebbian fashion with probability *A_ij_/S* proportional to its current strength (Fig. 2a, top); or (ii) with probability 1 – *p*, a random pair of neurons is selected for growth (Fig. 2a, bottom). In this way, the total connection strength *S* remains constant, with the wiring between neurons simply rearranging over time. Moreover, we note that the model only contains a single parameter *p*, representing the probability of Hebbian rather than random growth.

**Fig. 2.**
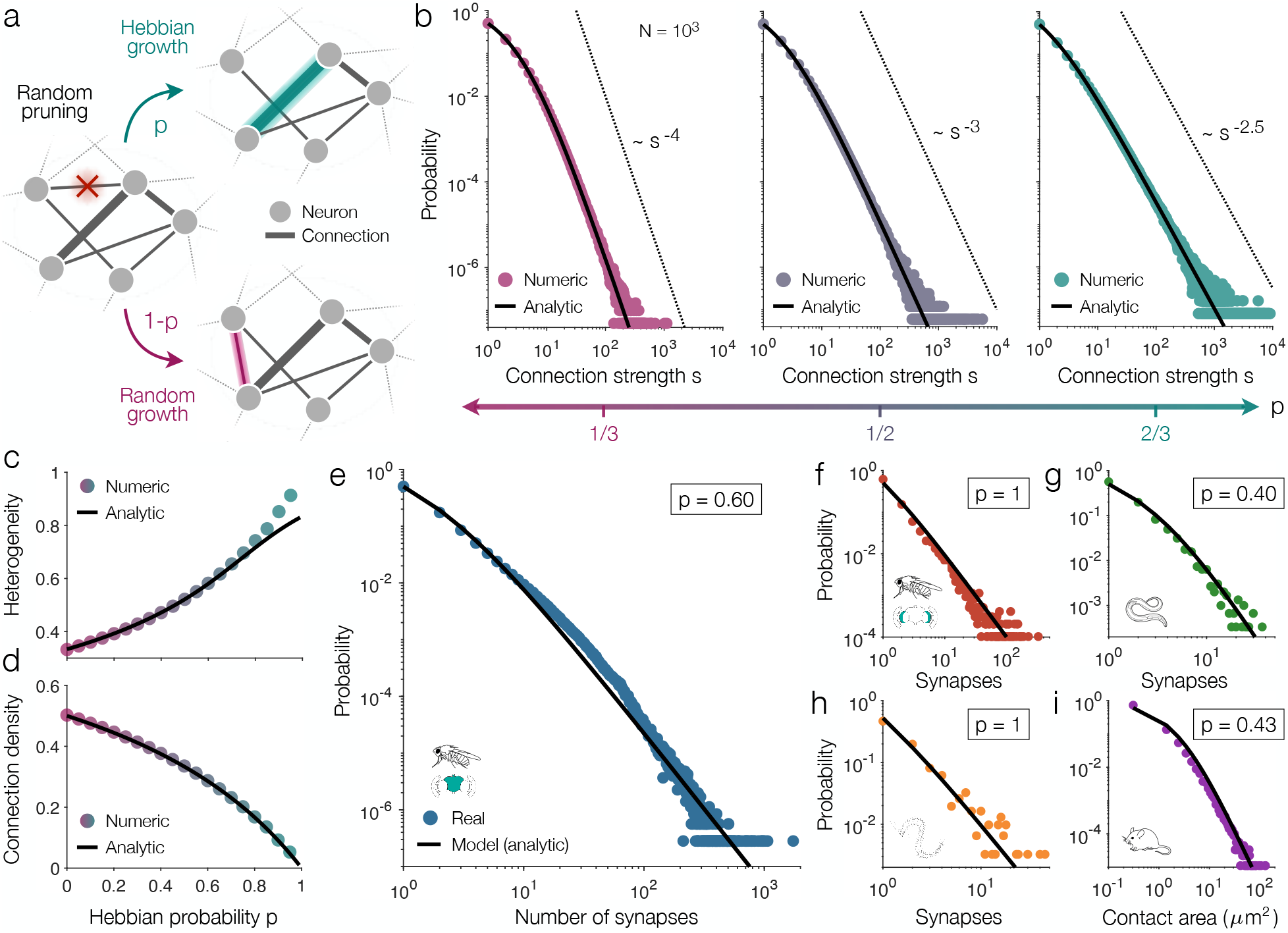
Hebbian dynamics produce power–law connection strengths. **a**, Minimal model of Hebbian network dynamics. At each time step, a random connection is pruned (left), and each unit of pruned strength is redistributed either in a Hebbian fashion (with probability *p*; top) or randomly (with probability 1 – *p*; bottom). **b**, Distributions of connection strengths for increasing values of the Hebbian probability *p* in networks of *N* = 10^3^ neurons and average connection strength 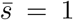. Data points depict numeric simulations (see Methods), solid lines represent the analytic prediction in Eq. (1), and dashed lines reflect power laws with the predicted exponent 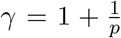. **c-d**, Connection heterogeneity (**c**) and density (**d**) as functions of the Hebbian probability *p*, computed using numeric simulations (data points) and the analytic distribution (lines). Heterogeneity 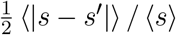 quantifies the heavy–tailedness of connectivity and is normalized to lie between zero and one,^37^ where 〈·〉 represents an average over *P*(*s*), and 〈|*s* – *s*′|〉 is the average absolute difference in connection strengths. Connection density is the fraction of all *N*(*N* – 1) possible connections with nonzero strength. **e-i**, Analytic model predictions from Eq. (1) with 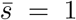 (to match the simulations in panels **b**-**d**) and *p* fit to the connectivity distributions (see Methods) of the *Drosophila* central brain (**e**),^1^ *Drosophila* optic medulla (**f**),^22^ *C. elegans* connectome (**g**),^28^ *Platynereis* sensory–motor circuit (**h**),^29^ and mouse retina (**i**).^21^

Ultimately, we wish to understand how the above dynamics impact the distribution *P*(*s*) of connection strengths *s*. Using the master equation method (see Methods), one can show that as the network evolves, the connection strengths approach an analytic steady–state distribution

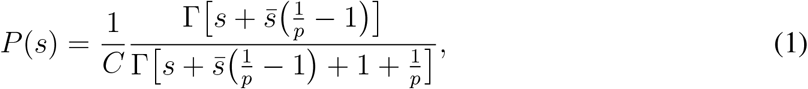

where 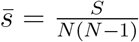 is the average connection strength, *C* is the normalization constant, and *Γ*(·) is Euler’s gamma function. Throughout the paper, we normalize all distributions to run over positive connection strengths *s* > 0 (that is, excluding null connections with *s* = 0). In the limit of strong connections 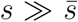, the above distribution falls off as a power law *P*(*s*) ~ *s*^-*γ*^, with a scale–free exponent 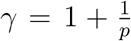 that depends only on the probability of Hebbian growth *p*. We confirm the analytic distribution in Eq. (1) and the power–law tail in numeric simulations (Fig. 2b).

As the proportion of Hebbian growth *p* decreases, the network becomes more random, limiting the heavy tail in *P*(*s*) and increasing the exponent *γ* (Fig. 2b, left). By contrast, if Hebbian growth dominates, the preferential attachment of synaptic strength yields a heavier tail and lower exponent *γ* (Fig. 2b, right). Indeed, as *p* increases, connection strengths become more heterogeneous (Fig. 2c) while the density of connections decreases (Fig. 2d), leading to the formation of sparse, heavy–tailed networks—precisely the features observed in real connectomes.^1,14,15,21,22,28,29,32^ We can even fit the Hebbian probability *p* (the only free parameter in the model) to the connectivity distributions of individual connectomes; for example, the *Drosophila* central brain is best described as arising from a combination of 60% Hebbian growth and 40% random growth (Fig. 2e). In fact, despite only fitting one parameter, our simple model provides a surprisingly good description of all the networks considered (Fig. 2e-i), with the proportion of Hebbian growth ranging from 40% for *C. elegans* (Fig. 2g) to 100% for the *Drosophila* optic medulla (Fig. 2f) and *Platynereis* sensory– motor circuit (Fig. 2h). Together, these results demonstrate that neurons can self–organize under minimal Hebbian dynamics to produce the kinds of sparse, heavy–tailed networks observed in real systems.

### Activity–dependent plasticity produces clustering

The model above generates networks with realistic connection strengths, even without including the effects of neuronal activity. Indeed, in classical Hebbian plasticity, synaptic connections evolve not just on their own, but also in response to the correlations (or functional connections) between neurons.^11,34–36^ By including this feedback between structural and functional connectivity, here we will show that additional properties of network structure emerge naturally. To introduce correlations between neurons, we must begin with a model of neuronal activity. While one can implement varying degrees of biological realism,^13,38^ for simplicity here we consider artificial neurons that, between each network update, reach steady–state activities *x_i_* defined by the self–consistent equations *x_i_* = tanh (*β* ∑_*j*_ *A_ij_x_i_*), where *β* ≥ 0 parameterizes the strength of interactions, and the connections *A_ij_* are implicitly normalized by 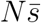 (see Methods). This model is equivalent to the mean–field Ising model,^39–41^ and incorporates standard nonlinear features of recurrent neural networks.^13,41,42^ The primary advantage of this model is the fact that, if the connections *A_ij_* are symmetric (as will be assumed in what follows), then we can use the fluctuation–dissipation theorem to derive a succinct analytic form for the covariances (see Methods): *C* = *βD*(*I* – *βDA*)^−1^, where 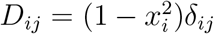 is a diagonal matrix and *I* is the identity.

We are now prepared to extend our network dynamics to include activity–dependent plasticity. The random pruning (Fig. 2a, left) and random growth (Fig. 2a, bottom) remain unchanged; but for Hebbian growth (Fig. 2a, top), rather than selecting a connection with probability proportional to its current strength *A_ij_*, we instead select a pair of neurons with probability proportional to their covariance *C_ij_* (see Methods). To gain intuition for this activity–dependent model, we can expand the covariances in the limit of weak interactions *β* ≪ 1, yielding

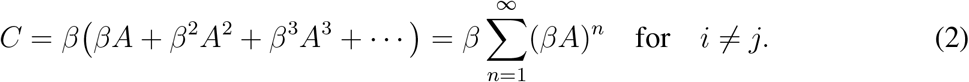

To lowest order in *β*, the covariances are proportional to the connection strengths *C* ~ *A*, and therefore Hebbian plasticity reinforces existing connections, exactly as in the original model (Fig. 3a, top). In this way, the original activity–independent dynamics (Fig. 2) can be viewed as the weak– interaction limit of the more general activity–dependent model. To second–lowest order in *β*, we have *C* ~ *A*^2^, and so Hebbian plasticity connects neurons that share common neighbors in the network (that is, neurons that interact via paths of length two). In this case, activity–dependent growth leads to the formation of triangles (Fig. 3a, middle), which, as we will see, produces clustering in the topology of connections. In general, as the interaction strength *β* increases toward one (the critical interaction strength for random networks; see Supplementary Information), Hebbian plasticity reinforces interactions of increasing topological distance, thereby producing longer and longer loops in the network (Fig. 3a, bottom).

**Fig. 3.**
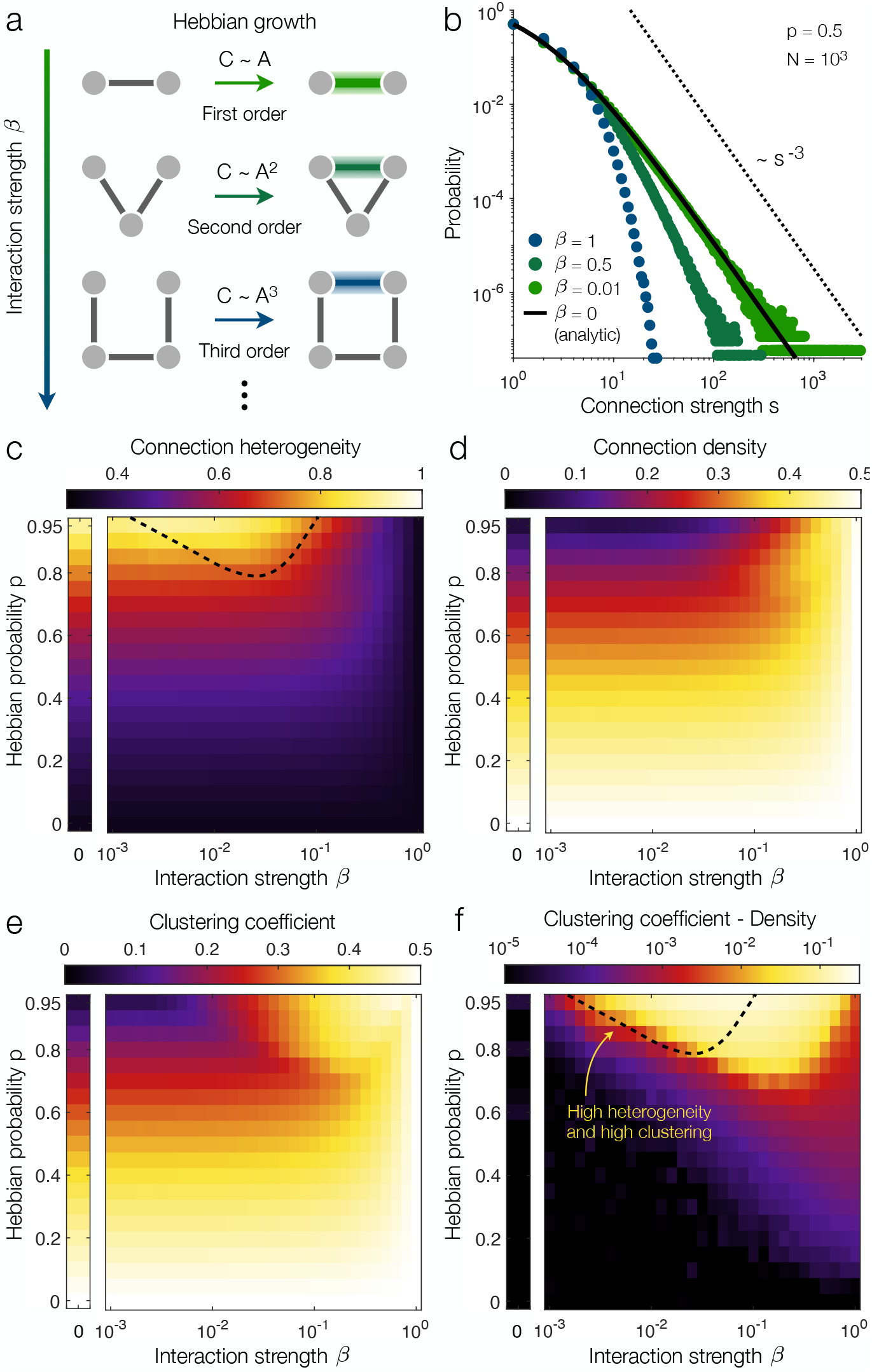
Activity–dependent dynamics give rise to network clustering. **a**, Expanding Hebbian growth in the limit of weak interactions. To lowest order in the interaction strength *β*, Hebbian growth reinforces existing connections (top); to second order, Hebbian growth reinforces interactions of length two, forming triangles (middle); in general, to *n*^th^ order, Hebbian growth reinforces interactions of length *n*, thus closing loops of length *n* + 1 (bottom). **b**, Distributions of connection strengths for different interaction strengths *β* in networks of *N* = 10^3^ neurons, Hebbian probability *p* = 0.5, and average connection strength 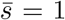. Solid line depicts the analytic activity–independent prediction in Eq. (1), and dashed line illustrates a power–law with exponent 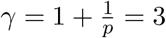. **c-f**, Phase diagrams of connection heterogeneity (**c**), connection density (**d**), clustering coefficient (**e**), and the difference between clustering coefficient and connection density (**f**) as functions of the Hebbian probability p and interaction strength *β* for simulated networks of *N* = 10^3^ neurons and average connection strength 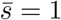 (see Methods). In each panel, for *β* = 0 (left), we plot values for the original activity–independent model (Fig. 2). In panels **c** and **f**, dashed lines indicate the region of parameter space with simultaneously heavy–tailed connection strengths and highly clustered topologies.

Simulating the above dynamics, we can examine how activity–dependent plasticity impacts both the strength and topology of connections. In the limit of weak interactions, because the activity–dependent and –independent dynamics become equivalent, the connection strengths converge to the heavy–tailed distribution in Eq. (1) (Fig. 3b). Moreover, since *C* ~ *A*, the model also produces heavy–tailed correlations, just as observed in large–scale recordings of neuronal activity (Fig. 1h; see Supplementary Information). By contrast, as *β* increases, Hebbian growth fails to reinforce existing connections (Fig. 3a), thereby constraining the heavy tail of connection strengths (Fig. 3b). Indeed, we find that the heterogeneity of connection strengths increases with the Hebbian probability *p* and decreases with the interaction strength *β* (Fig. 3c). Conversely, the density of connections decreases with p and increases with *β* (Fig. 3d), such that sparse, heavy–tailed connectivity arises for large Hebbian probabilities and weak to intermediate interactions.

While to first order, activity–dependent plasticity yields heavy–tailed connectivity, as discussed above, the second–order effect is the formation of triangles (Fig. 3a). Notably, real neuronal networks are known to exhibit a high density of triangles, a property known as clustering.^6–9^ Clustering is typically quantified using the ratio *N*_Δ_/*N*_^_ of the number of neuron triangles *N*_Δ_ to the number of triplets *N*_^_ (sets of three neurons with at least two connections). For random networks, this clustering coefficient is identical to the density of connections; and in fact, we see that the clustering coefficient (Fig. 3e) closely tracks the connection density (Fig. 3d) across most values of *p* and *β*. In particular, as interactions vanish (*β* → 0), the original activity–independent dynamics (Fig. 2) yield random network topologies (for all values of *p*), such that the clustering coefficient becomes identical to the connection density (Fig. 3f, left). Thus, if interactions are too strong, then connection strengths lose their heavy tail (Fig. 3b,c), and if interactions are too weak, then the network fails to develop clustering (Fig. 3f). Yet for intermediate interaction strengths *β* (and large Hebbian probabilities *p*), the activity–dependent dynamics produce both heavy–tailed connection strengths and highly clustered topologies (Fig. 3c,f, highlighted regions), two fundamental features of real neuronal networks.^1–9^

The activity–dependent dynamics qualitatively generate realistic network structures, but can they quantitatively reproduce the properties of specific neuronal networks? For example, in the *Drosophila* central brain (Fig. 1a-b), we find that the connections are more heterogeneous (Fig. 4a) and more clustered (Fig. 4b) than comparable random networks; and we confirm the same results across all of the connectomes analyzed in Fig. 1 (see Supplementary Information). Simulating networks with the same average connection strength 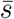 as the *Drosophila* central brain, the original activity–independent model (Fig. 2) can capture the connection heterogeneity observed in the data (Fig. 4a), but (due to its random topology) fails to produce realistic clustering (Fig. 4b). However, by generalizing to activity–dependent plasticity, the networks can simultaneously attain both the connection heterogeneity and clustering observed in the *Drosophila* connectome (Fig. 4c). In fact, by fitting only two parameters (the Hebbian probability *p* and interaction strength *β*), our activity– dependent model quantitatively reproduces the connection heterogeneity and clustering in all of the connectomes analyzed (Fig. 4d). Furthermore, we find that the activity–dependent dynamics capture additional features of neuronal connectivity, such as the relationship between connection strength and clustering (see Supplementary Information). Thus, by extending the network dynamics to include activity–dependent plasticity, the connections between neurons self–organize to produce not only the heavy–tailed connectivity, but also the clustering observed in real networks.

**Fig. 4.**
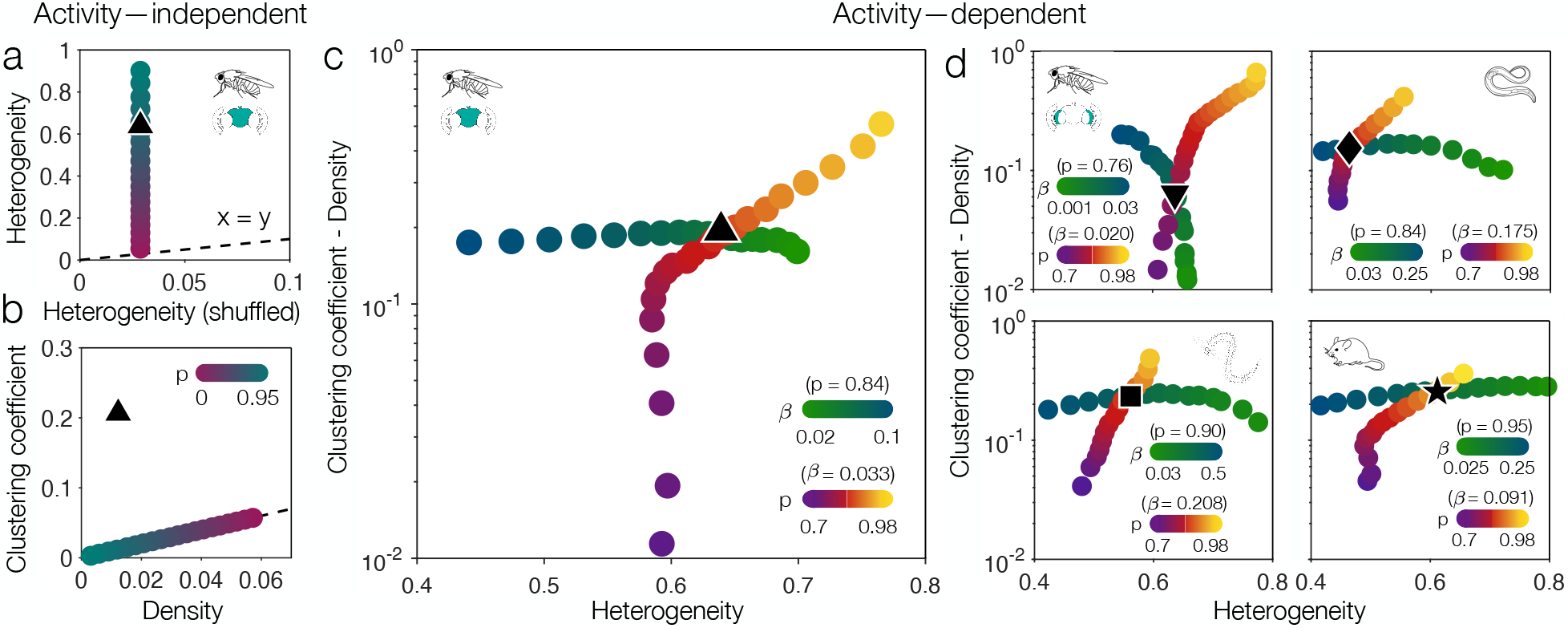
Capturing heavy–tailed and clustered connectivity in real connectomes. **a**, Heterogeneity of connection strengths in the *Drosophila* central brain before and after shuffling the synapses among the neurons. Colored points depict the activity–independent model while sweeping over the Hebbian probability *p* for networks of *N* = 10^3^ neurons and average connection strength 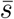 matching the *Drosophila* central brain (see Methods). **b**, Clustering coefficient versus connection density for the same networks as panel **a**. **c**, Clustering coefficient minus connection density (that is, the clustering beyond that of a network with randomized connections) versus connection heterogeneity. Colored points illustrate the activity–dependent model (with *N* = 10^3^ and 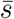 matching the *Drosophila* central brain) while sweeping over the interaction strength *β* (blue/green) or Hebbian probability *p* (purple/yellow). For *p* = 0.84 and *β* = 0.033, the activity–dependent dynamics yield the same connection heterogeneity and clustering coefficient (minus density) as the *Drosophila* central brain. **d**, Same as panel **c**, but for the *Drosophila* optic medulla (top left),^22^ *C. elegans* connectome (top right),^28^ *Platynereis* sensory–motor circuit (bottom left),^29^ and mouse retina (bottom right).^21^ Across all panels, to compute clustering and compare against model networks, the real connectomes are symmetrized, such that the connectivity is defined by *A* + *A^T^* (see Methods). In panels **c-d**, to improve sampling, data points for the activity–dependent model are averaged across adjacent parameter values.

## Discussion

Across a diverse range of species, the strengths of neuronal connections are similarly heavy– tailed (Fig. 1), suggesting a common underlying mechanism. Here, we develop a minimal model in which synaptic connectivity self–organizes under a mixture of Hebbian and random dynamics. The resulting networks exhibit scale–free distributions of connectivity strength, with an exponent 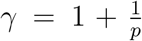 that depends only on the proportion *p* of Hebbian growth (Fig. 2). Moreover, by generalizing the model to include activity–dependent plasticity, the network dynamics naturally give rise to clustering (Fig. 3), another prominent feature of neuronal connectivity (Fig. 4). Given the simplicity of the model, future work can immediately begin to include more realistic plasticity mechanisms, which, in turn, may produce additional properties observed in neuronal connectomes. For example, rather than updating the synaptic connectivity based on equal–time correlations, one could further generalize the activity–dependent model to include spike–time–dependent plasticity, wherein synapses are updated based on time–delayed correlations.^11,35^ These temporal asymmetries will naturally generate asymmetries in the connections themselves, a central property of real connectomes.^1–7,14,15,21,22,28,29^ Additionally, one can directly implement more biophysically realistic models of neuronal activity.38 We remark, however, that as long as the correlations between neurons decrease with distance in the network, one should still expect activity–dependent plasticity to produce clustering, just as in our model (Fig. 3). Together, the results and models developed here demonstrate how two fundamental features of connectomes—heavy–tailed connectivity and clustering—can arise from simple network dynamics, providing a framework for future investigations into the self–organization of neuronal connectivity.

## Methods

### Neuronal connectivity data

The datasets analyzed in this paper were each previously published and are openly accessible.^1,21,22,28,29,33^ All connectomes were sectioned and imaged at the neuron level using electron microscopy.^1,21,22,28,29^ The neurons and synapses (or contacts for the mouse retina) were then traced using a mixture of manual labelling and automated image processing. For further experimental details concerning the *Drosophila* central brain (Fig. 1a-b),^1^ *Drosophila* optic medulla (Fig. 1c),^22^ *C. elegans* (Fig. 1d),^28,43^ *Platynereis* sensory–motor circuit (Fig. 1e),^29^ and mouse retina (Fig. 1f-g),^21^ see the respective references. We have made the lists of neurons and connections between them openly accessible in a common format (see Data Availabaility).

When comparing against the activity–independent model (Figs. 1 and 2), the connections *A_ij_* are directed, reflecting the number of synapses (or, for the mouse retina, the number or area of contacts) from one neuron *i* to another *j*. When comparing against the activity–dependent model (Fig. 4), we use symmetrized versions of the connectomes, where the connections *A* + *A^T^* reflect the number of synapses (or contact area) between neurons in both directions. In Table 1, we list the network properties for both directed and symmetrized versions of the different connectomes.

**Table 1.** Network properties of neuronal connectomes. For each connectome, we list the number of neurons *N*, total number of synapses (or contact area for the mouse retina) *S*, average connection strength 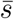, number of nonzero connections, connection heterogeneity, connection density, and clustering coefficient. The values outside (inside) parentheses correspond to directed (symmetrized) versions of the connectomes.

|  | *Drosophila*<br>central brain | *Drosophila*<br>optic medulla | *C. elegans*<br>connectome | *Platynereis*<br>sensory-motor | Mouse<br>retina |
| --- | --- | --- | --- | --- | --- |
| $N$ | 21,739 | 1,781 | 279 | 79 | 1,123 |
| $S$ | $1.43 \times 10^7$ | $3.35 \times 10^4$ | $6.82 \times 10^3$ | $1.09 \times 10^3$ | $9.67 \times 10^4 \mu m^2$ |
| $\bar{s}$ | 0.030 | 0.011 | 0.088 | 0.177 | $0.077 \mu m^2$ |
| # connections | $3.55 (2.90) \times 10^6$ | $9.63 (8.91) \times 10^3$ | $2.99 (2.29) \times 10^3$ | 300 (270) | $9.00 (9.00) \times 10^4$ |
| Heterogeneity | 0.616 (0.639) | 0.626 (0.637) | 0.433 (0.463) | 0.556 (0.562) | 0.612 (0.612) |
| Density | 0.008 (0.012) | 0.003 (0.006) | 0.039 (0.059) | 0.049 (0.088) | 0.071 (0.143) |
| Clustering coeff. | (0.207) | (0.069) | (0.214) | (0.321) | (0.398) |

To investigate covariances in neuronal activity (Fig. 1h and Supplementary Information), we study a previously– published dataset of two–photon calcium recordings of large neuronal populations in the visual cortex of awake mice.^33^ To account for variations in fluorescence across neurons, we binarize the preprocessed activity traces to reflect, at each moment in time, whether a neuron is above or below its average activity. For further details on experimental setup and data processing, see the reference herein.^33^

### Simulating the activity–independent model

In the activity–independent model (Fig. 2), we begin with *N* neurons, *S* = ∑_*ij*_ *A_ij_* units of connection strength (for an average connection strength of 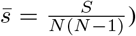 and a probability *p* of Hebbian growth. The network then evolves as described in the main text. Throughout the paper, we simulate networks with *N* = 10^3^ neurons, for a total of *N*(*N* – 1) ≈ 10^6^ possible directed connections, and (unless otherwise specified) an average connection strength of 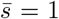. We initialize each network by distributing the strength *S* randomly among the possible connections. When computing distributions (as in Fig. 2b) or quantities of interest (as in Fig. 2c-d), we take 100 network samples; between each sample, we prune *N*(*N* – 1) different connections, such that each directed pair of neurons is pruned once on average. We also prune each connection an average of 50 times for burn–in before taking any samples. In practice, to speed up calculations, one can run multiple network updates in parallel; here we prune 1% of the possible connections at a time, and redistribute all of the pruned strength simultaneously. We confirm that this parallelization does not significantly affect the accuracy of the simulations (Fig. 2b). In Figs. 3 and 4, to compare against the activity–dependent model, we study symmetric (as opposed to directed) networks; in practice, the only difference for the simulations is that there are *N*(*N* – 1)/2 possible connections (neurons pairs) rather than *N*(*N* – 1) directed pairs, such that the total synaptic strength is given by 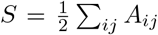. Code for simulating the activity–independent model has been made openly available (see Code Availability).

### Deriving the distribution of connection strengths

As a network evolves according to the activity–independent dynamics, the connection strengths converge to the steady–state distribution *P*(*s*) defined in Eq. (1). To derive *P*(*s*), we begin by writing down the master equation describing the change 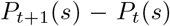 in the connection strength distribution from one time step of the dynamics *t* to the next. During each network update, a random connection is pruned, yielding an average decrease in probability of 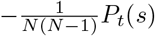. On average, the pruning removes 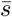 units of connection strength. For each unit of pruned strength, we perform Hebbian growth with probability *p*, selecting each connection of strength *s* with probability *s/S*. On average, this Hebbian growth yields an increase in probability of 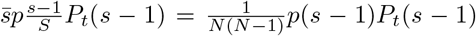 and a decrease of 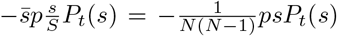. Finally, for each unit of pruned strength, we perform random growth with probability 1 – *p*, yielding (on average) an increase in probability of 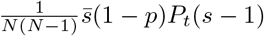 and a decrease of 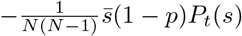.

Combining the above contributions, we arrive at the following master equation,

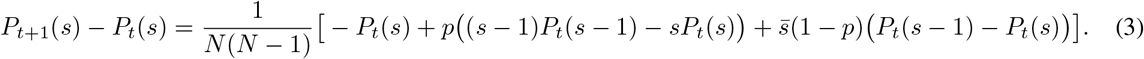

As the network evolves, the connection strengths approach a steady–state distribution *P*_*t*+1_(*s*) = *P_t_*(*s*) = *P*(*s*), which transforms Eq. (3) into the following recursive relation,

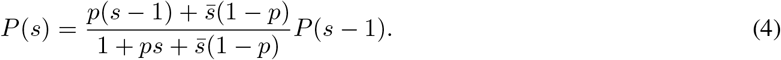

Solving for *P*(*s*), we arrive at the distribution in Eq. (1), which we note is normalized throughout the paper to run over positive connection strengths *s* > 0 (that is, excluding connections of strength zero). In the limit of strong connections 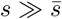, one can show that *P*(*s*) scales as *s*^-*γ*^, where 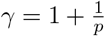. The analytic steady–state distribution in Eq. (1) and the power–law tail are confirmed numerically in Fig. 2b. To compute the Hebbian probability *p* that best describes a given connectome (as in Fig. 2e-i), we run gradient descent [using the analytic distribution in Eq. (1)] on the root–mean–square error of the log probabilities.

### Covariances in neuronal activity

To study activity–dependent plasticity, we implement model neurons that, between each network update, reach steady–state activities *x_i_* defined by the nonlinear equation *x* = tanh[*β*(*Ax* + *b*)], where *β* is the interaction strength, *b_i_* are the biases (here assumed to be zero for simplicity), and tanh(·) is applied component–wise. Additionally, as is common, we implicitly normalize the connections *A_ij_* by 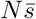, such that the total interaction strength per neuron approaches one for large networks. This system is equivalent to a mean–field Ising model with inverse temperature *β*, interactions *A_ij_*, and external fields *b_i_* = 0. As such, for symmetric connections *A* = *A^T^*, one can use the fluctuation–dissipation theorem to compute the covariances in activity,

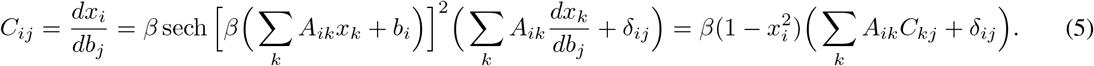

Given a stable solution *x* for the network activity, one can solve for the covariances *C* = *βD*(*I* – β*DA*)^-1^, where 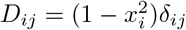 and *I* is the identity.

The above expression for the covariances holds for arbitrary interaction strengths *β* and biases *b*. Yet for zero biases *b* = 0, things simplify even further: for sufficiently weak interactions *β* < *ρ*(*A*), where *ρ*(·) is the spectral radius (or the magnitude of the largest eigenvalue), the unique solution to the self–consistent equation is *x* = 0. In this case, we have *D* = *I*, yielding the weak–interaction expansion

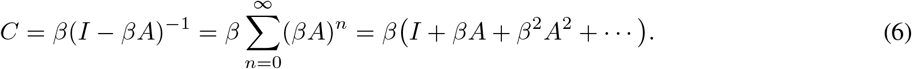

As the interaction strength *β* approaches *ρ*(*A*), the solution *x* = 0 becomes unstable, and the covariances in Eq. (6) diverge;^40^ in this sense, *ρ*(*A*) defines the critical interaction strength. For example, if the connection strength *S* is distributed randomly, then one can show that *ρ*(*A*) = 1 in the limit *N* → ∞ (see Supplementary Information). Thus, for random connectivity, the weak–interaction expansion only holds for *β* < 1.

### Simulating the activity–dependent model

In the activity–dependent model, the network evolves precisely as in the activity–independent model (with symmetric connections), except for one key difference: during Hebbian growth, rather than selecting a connection with probability proportional to its strength *A_ij_*, we select a pair of neurons with probability proportional to their covariance *C_ij_* (we note that the pair of neurons need not be connected previously). Thus, prior to each iteration of the dynamics, we must compute the covariances *C*, as described above. This, in turn, requires computing a solution *x* for the average activities of the neurons, which can be accomplished efficiently by initializing *x* = –1 and iterating the nonlinear equation *x* ← tanh(*βAx*) until convergence.^40^ In simulations, we consider networks with *N* = 10^3^ neurons, for a total of *N*(*N* – 1)/2 ≈ 5 × 10^5^ possible undirected connections, and (unless otherwise specified) an average connection strength 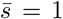. As in the activity–independent model, for each calculation we record 100 samples, between which each possible connection is pruned once on average, and before which each possible connection is pruned 50 times for burn–in. In the phase diagrams in Fig. 3c-f, for *p* ≥ 0.7, in order to improve sampling we average over ten simulations for each data point. Code for simulating the activity–dependent model has been made openly available (see Code Availability).

## Data Availability

The neuronal data analyzed in this paper are openly available at github.com/ChrisWLynn/Heavy_tailed_connectivity.

## Code Availability

The code used to perform the analyses in this paper is openly available at github.com/ChrisWLynn/Heavy_tailed_connectivity.

## Supplementary Information

Supplementary text and figures accompany this paper.

## Acknowledgements

The authors thank David J. Schwab and Lia Papadopoulos for discussions and feedback and Paschalis Kratsios for help with illustrations. This work was supported in part by the National Science Foundation, through the Center for the Physics of Biological Function (PHY–1734030) and a Graduate Research Fellowship (C.M.H.); by the James S. McDonnell Foundation through a Postdoctoral Fellowship Award (C.W.L.); and by the National Institutes of Health BRAIN initiative (R01EB026943).

## Citation diversity statement

Recent work in several fields of science,^44–48^ and physics in particular,^49^ has identified citation bias negatively impacting women and other minorities. Here we sought to proactively consider choosing references that reflect the diversity of the field in thought, form of contribution, gender, and other factors. Excluding (including) self-citations to the current authors, our references contain 43% (41%) women lead authors and 36% (39%) women senior authors.

## Author Contributions

C.W.L. and S.E.P. conceived the project. C.W.L. designed the models, and C.W.L. and C.M.H. performed the analysis with input from S.E.P. C.W.L. wrote the manuscript and Supplementary Information; and C.M.H. and S.E.P. edited the manuscript and Supplementary Information.

