## Supplementary Information for "Heavy–tailed neuronal connectivity arises from Hebbian self–organization"

#### Contents

**1 Introduction** **2**

**2 Heavy–tailed correlations in mouse visual cortex** **2**

**3 Critical interaction strength in random networks** **4**

**4 Activity–dependent model generates heavy–tailed correlations** **4**

**5 Connection heterogeneity and clustering in real connectomes** **6**

**6 Predicting the dependence of clustering on connection strength** **7**

**References** **9**

### 1 Introduction

In this Supplementary Information, we provide extended analysis and discussion to support the results presented in the main text. The sections are ordered to align with their references in the main text. In Sec. 2, we explore the distributions of covariances in activity of neurons in the visual cortex of mice responding to different stimuli. Notably, across multiple mice and distinct stimuli, we find that the covariances are heavy–tailed, similar to the structural connections explored in the main text. In Sec. 3, for networks with random connectivity, we prove that the critical interaction strength in the mean–field Ising model is $\beta = 1$. In Sec. 4, we show that our activity–dependent model of network dynamics gives rise not only to heavy–tailed distributions of structural (i.e., synaptic) connections, but also to heavy–tailed distributions of functional (i.e., correlational) connections, just as observed in real data (see Sec. 2). In Sec. 5, we show that the connectomes analyzed in the main text have more heterogeneous connection strengths and more clustered topologies than comparable random networks. Finally, in Sec. 6, we show that the activity–dependent model predicts the relationship between connection strength and clustering observed in real connectome data.

### 2 Heavy–tailed correlations in mouse visual cortex

Throughout the main text, we focus on "structural" neuronal networks, reflecting the physical wiring between cells. One could additionally consider networks of "functional" connections, defined by the correlations in activity between neurons. Investigating the large–scale properties of functional networks is now possible due to advances in neuroimaging, which have enabled simultaneous activity recordings of large neuronal populations in vivo. Here, we consider two–photon calcium recordings of over $10,000$ neurons in the mouse visual cortex, measured and described in previously–published work.[1] In Fig. 1h in the main text, for one mouse responding to natural images, we show that the distribution of covariances between neurons is heavy–tailed, following a similar form to the distributions of structural connections observed across different animals (Fig. 1b-e,g in the main text). Here, we show that these heavy–tailed correlations are not limited to a specific mouse, nor to natural images as a stimulus, but instead appear to be a general feature

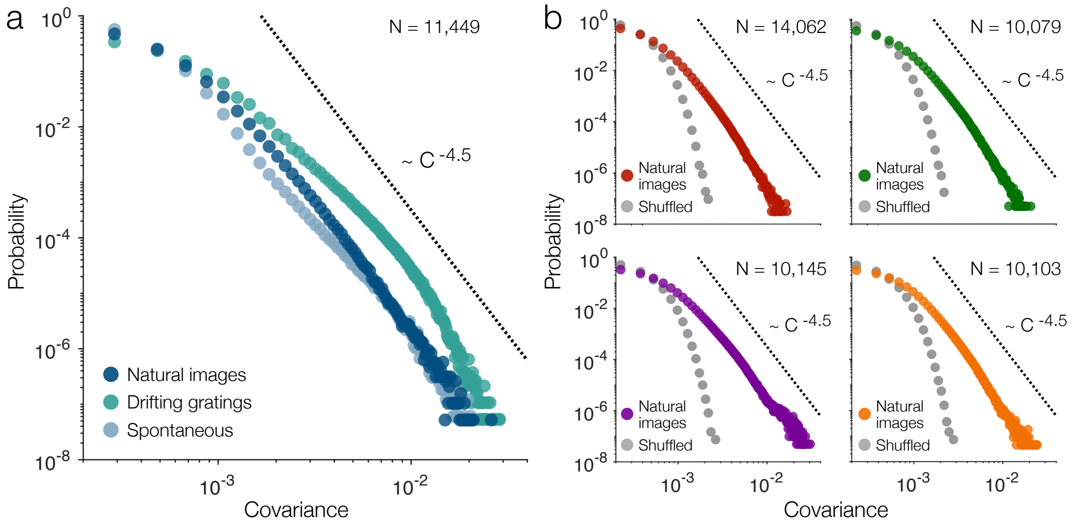


**Fig. S1 | Heavy–tailed covariances in large neuronal populations.** **a**, Distributions of covariances in neuronal activity (recorded using two–photon calcium imaging) in the mouse visual cortex while responding to different stimuli.[1] **b**, Distributions of covariances in the visual cortices of different mice while responding to natural images (colors) and after shuffling the activity time–series (grey).

of large–scale neuronal activity.

In Fig. S1a, we show the distribution of covariances between neurons in the same mouse as Fig. 1h in the main text, but for three different stimuli: (i) natural images, (ii) a drifting grating of dark lines, and (iii) no stimulus (spontaneous activity). Across all three stimuli, we find that the covariances follow similar heavy–tailed distributions, with the vast majority of neurons behaving nearly independently while a select few pairs are strongly correlated. Additionally, for different mice all responding to the same set of natural images, we find that the distributions of covariances are similarly heavy–tailed (Fig. S1b). By contrast, after removing the statistical dependencies between neurons by shuffling their activities in time, we find that the heavy tails of strong correlations are destroyed (Fig. S1b, grey points). Together, these results suggest that heavy–tailed functional connectivity (or correlations) may be a general feature of neuronal activity. In Sec. 4, we show that precisely these types of heavy–tailed correlations emerge naturally from our activity–dependent model.

#### 3 Critical interaction strength in random networks

In the main text, we study activity–dependent plasticity using model neurons with average activities given by the self–consistent equation $x = \tanh(\beta \tilde{A} x)$, where $x_i$ is the average activity of neuron $i$, $\beta$ is the interaction strength, and $\tilde{A}_{ij} = \frac{1}{N\bar{s}} A_{ij}$ is the normalized connection strength from $j$ to $i$ (where $A_{ij}$ is the adjacency matrix of the network, $N$ is the size of the network, and $\bar{s} = \frac{1}{N(N-1)} \sum_{ij} A_{ij}$ is the average connection strength). This system has a critical interaction strength of $\beta = \rho(\tilde{A}) = \frac{1}{N\bar{s}} \rho(A)$, where $\rho(\cdot)$ is the spectral radius (or magnitude of the largest eigenvalue). For $\beta < \rho(\tilde{A})$, the unique solution for the neuronal activities is $x = 0$, and one can derive a weak–interaction expansion for the covariances between neurons (see Methods in the main text). As $\beta$ approaches $\rho(\tilde{A})$, the solution $x = 0$ becomes unstable and the covariances diverge.[2]

To guide intuition, here we show that, if connectivity is placed randomly in the network, then the critical interaction strength approaches $\rho(\tilde{A}) = 1$ in the thermodynamic limit $N \to \infty$. To begin, it is know that for a nonnegative matrix $A$, the spectral radius falls between the minimum and maximum row sums, such that

$$\min_i \sum_j A_{ij} \leq \rho(A) \leq \max_i \sum_j A_{ij}. \tag{S1}$$

If each unit of connection strength is placed between a random pair of neurons, then the connection strengths $A_{ij}$ follow a Poisson distribution with mean and variance $\bar{s}$. Since each connection strength $A_{ij}$ is Poisson–distributed, each row sum $\sum_j A_{ij}$ also follows a Poisson distribution with mean and variance $N\bar{s}$. Thus, the normalized row sums $\sum_j \tilde{A}_{ij}$ have mean one and variance $\frac{1}{N\bar{s}}$. Combined with Eq. (S1), this establishes that the critical interaction strength $\rho(\tilde{A})$ tends to one as $N \to \infty$.

#### 4 Activity–dependent model generates heavy–tailed correlations

In Fig. S1 (and Fig. 1h in the main text), we show that covariances in neuronal activity can be heavy–tailed. Here, we will show that such heavy–tailed correlations arise naturally in our activity–dependent model.

In the main text, we showed that the correlations $C_{ij}$ are related to the connection strengths

$A_{ij}$ via the expression

$$C = \beta D(I - \beta D\tilde{A})^{-1}, \tag{S2}$$

where $I$ is the identity, $\tilde{A} = \frac{1}{N\bar{s}}A$ is the normalized connectivity matrix, $D_{ij} = (1 - x_i^2)\delta_{ij}$ is a diagonal matrix, and $x$ is a stable solution to the self–consistency equation. In the weak–interaction limit $\beta \ll 1$, the only stable solution is $x = 0$, and so the covariances simplify significantly,

$$C = \beta(I - \beta\tilde{A})^{-1} = \beta\sum_{n=0}^{\infty}(\beta\tilde{A})^n = \beta\big(I + \beta\tilde{A} + \beta^2\tilde{A}^2 + \beta^3\tilde{A}^3 + \dots\big). \tag{S3}$$

For the off-diagonal entries in Eq. (S3)—that is, for the correlations between neurons—to lowest order in $\beta$, we have $C = \beta^2\tilde{A} = \frac{\beta^2}{N\bar{s}}A$. Thus, for weak interactions, we should expect the covariances $C_{ij}$ to have the same distributional form as the connection strengths $A_{ij}$; specifically, the distribution of covariances should have a heavy tail that drops off as a power law with exponent $\gamma = 1 + \frac{1}{p}$. As interactions become stronger, the covariances $C_{ij}$ should include progressively longer–range connections between $i$ and $j$, until, at the critical interaction strength $\beta = \frac{1}{N\bar{s}}\rho(A)$, all neurons should become strongly correlated.

In Fig. S2, we plot the distributions of the normalized covariances $\frac{N\bar{s}}{\beta^2}C_{ij}$. As discussed above, in the weak–interaction limit $\beta \to 0$, these normalized covariances should equal the connectivity strengths $A_{ij}$, which, as discussed in the main text, are distributed according to the analytic form

$$P(s) = \frac{1}{C}\frac{\Gamma\big[s + \bar{s}\big(\frac{1}{p} - 1\big)\big]}{\Gamma\big[s + \bar{s}\big(\frac{1}{p} - 1\big) + 1 + \frac{1}{p}\big]}, \tag{S4}$$

where

$$C = \sum_{s=1}^{\infty}\frac{\Gamma\big[s + \bar{s}\big(\frac{1}{p} - 1\big)\big]}{\Gamma\big[s + \bar{s}\big(\frac{1}{p} - 1\big) + 1 + \frac{1}{p}\big]} \tag{S5}$$

is the normalization constant. Indeed, as the the interaction strength $\beta$ decreases toward zero, we see that the distribution of normalized covariances approaches Eq. (S4), with a heavy tail that drops off as a power law, as predicted (Fig. S2, orange). As $\beta$ increases, however, the covariances begin to lose their heavy tail (Fig. S2, red). As the interaction strength $\beta$ approaches one (the critical interaction strength for random networks; see Sec. 3), the distribution of correlations loses its heavy tail entirely, with all neurons becoming strongly correlated with one another (Fig. S2,

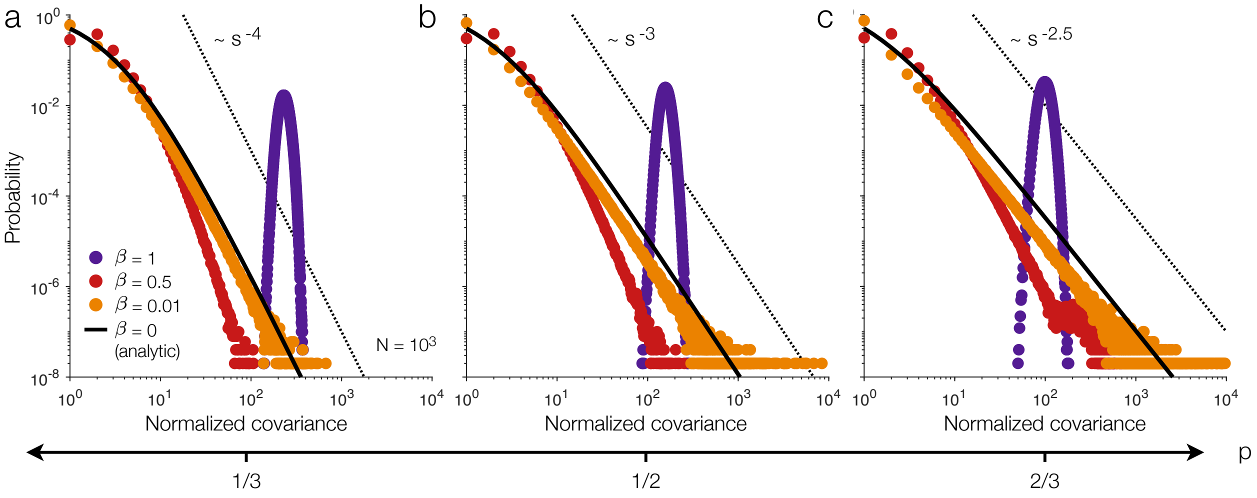


**Fig. S2 | Heavy–tailed correlations arise from activity–dependent plasticity.** Distributions of normalized covariances $\frac{N\bar{s}}{\beta^2}C_{ij}$ generated by the activity–dependent model for Hebbian probabilities $p = 1/3$ (**a**), $p = 1/2$ (**b**), and $p = 2/3$ (**c**). Solid lines illustrate the analytic connectivity strength distribution in Eq. (S4), and dashed lines reflect power laws with the predicted exponent $\gamma = 1 + \frac{1}{p}$. Data points depict simulations of networks with $N = 10^3$ neurons with average connectivity strength $\bar{s} = 1$.

purple). Notably, these results hold across different values of the Hebbian probability $p$ (Fig. S2). Thus, for intermediate and weak interactions, our activity–dependent model generates precisely the types of heavy–tailed correlations observed in large-scale neuronal recordings (Fig. 1h in the main text and Fig. S1).

#### 5 Connection heterogeneity and clustering in real connectomes

In the main text, we show that the *Drosophila* central brain connectome has larger connection heterogeneity and clustering than comparable random networks (Fig. 4a-b in the main text). Here, we show that these results also hold for the other networks analyzed in the paper. In Fig. S3a, we plot the connection heterogeneity of the different connectomes against the heterogeneity of equivalent networks with the synapses (or contacts for the mouse retina) shuffled. Similarly, in Fig. S3b, we plot the clustering coefficient of the real connectomes versus the density of connections, which is equivalent to the clustering coefficient of networks with the connections (as opposed to the synapses) shuffled. To be clear, the randomized networks in Fig. S3a have the total connection strength (i.e., number of synapses or connection area) held fixed, while the connection density in

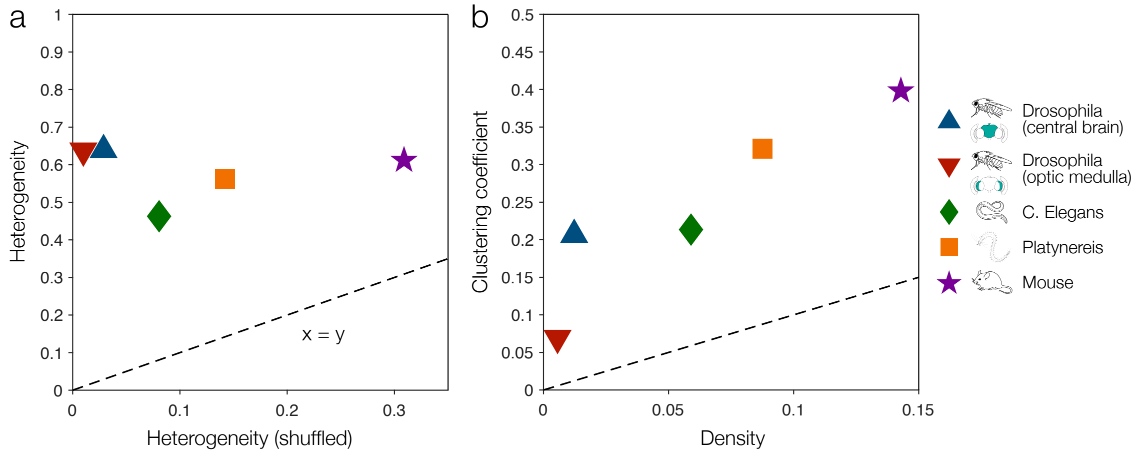


**Fig. S3 | Network properties of neuronal connectomes. a**, Connection heterogeneity of the connectomes analyzed in the main text versus that of randomized networks with the synapses (or contacts for the mouse retina) shuffled between the neurons. **b**, Clustering coefficient versus connection density for the real networks. Connection heterogeneity is given by $\frac{1}{2}\langle |s - s'| \rangle \,/\, \langle s \rangle$, where $\langle \cdot \rangle$ denotes an average over non–zero connection strengths. Clustering coefficient is the ratio $N_\triangle / N_\wedge$, where $N_\wedge$ is the number of neuron triplets with at least two connections, and $N_\triangle$ is the number of neuron triangles (that is, the number of neuron triplets with three connections). Connection density is the fraction of neuron pairs with a connection.

Fig. S3b is equivalent to the clustering coefficient for randomized networks with the number of connections held fixed. In both panels, we see that all of the data points lie above the $x = y$ line, indicating that the real networks have more heterogeneous connections and more clustered topologies than one would expect for comparable random networks.

#### 6 Predicting the dependence of clustering on connection strength

We have shown that the activity–dependent model is capable of quantitatively reproducing the heavy–tailed connection strengths and clustering observed in real connectomes (Figs. 3 and 4 in the main text). Here we show that the activity–dependent model can also predict the *relationship* between connection strength and clustering. By thresholding a network to only include connections above a desired strength, one can investigate how clustering (or other network properties) vary as we focus on stronger and stronger connections. In the *Drosophila* central brain connectome,[3] as we increase the threshold, the density of connections decreases (as expected), yet the clustering coefficient remains relatively constant (Fig. S4, blue). By contrast, after shuffling the synapses

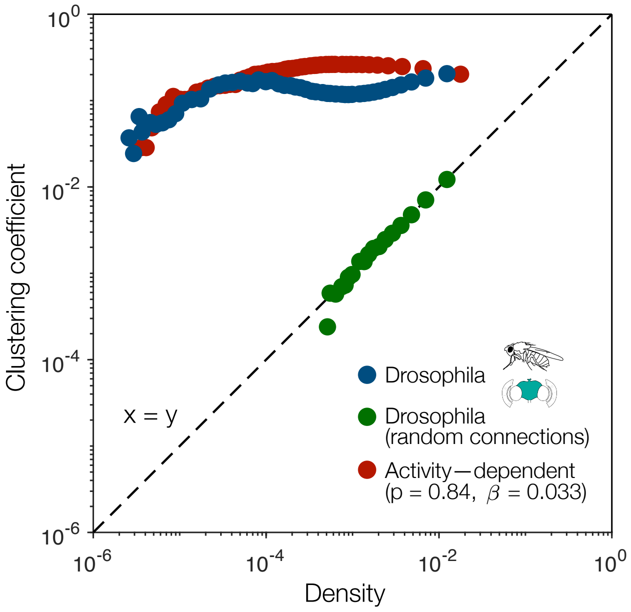


**Fig. S4 | Predicting the dependence of clustering on connection strength.** Clustering coefficient versus connection density for thresholded versions of the *Drosophila* central brain[3] before (blue) and after (green) shuffling the synapses among the neurons, and for the activity–dependent model with Hebbian probability $p = 0.84$ and interaction strength $\beta = 0.033$ (red). For the activity–dependent model (red), we simulate networks with $N = 10^3$ neurons and an average connection strength $\bar{s}$ matching that of the *Drosophila* central brain. For the *Drosophiphila* central brain (blue and green), to match the simulations of the activity–dependent model, we sample 100 random subnetworks of $N = 10^3$ neurons each.

between neurons, the dependence of clustering on connection strength is destroyed, and the clustering coefficient simply tracks the connection density Fig. S4, green). Finally, we can perform the same analysis on networks simulated using our activity–dependent model, with parameters $p = 0.84$ and $\beta = 0.033$ chosen to match the connection heterogeneity and the clustering coefficient (minus connection density) of the *Drosophila* central brain (see Fig. 4 in the main text). As we threshold to progressively stronger connections, we find that the simulated networks closely match the clustering of the real connectome (Fig. S4, red). Thus, our activity–dependent model is capable not only of reproducing the heavy–tailed connection strengths and clustering observed in real networks (Fig. 4 in the main text), but also the relationship between the two.
